# Context-independent expression of spatial code in hippocampus

**DOI:** 10.1101/2022.03.28.486068

**Authors:** S. Kapl, F. Tichanek, F. Zitricky, K. Jezek

**Affiliations:** Faculty of Medicine in Pilsen, Charles University, Pilsen, 32300 Czech Republic

**Keywords:** hippocampus, spatial representation, place cell, episodic memory, attractor, theta oscillations

## Abstract

Hippocampus plays a crucial role in formation and retrieval of spatial memory across mammals and episodic memory in humans. Episodic as well as spatial memories can be retrieved irrespectively of subject’s awake behavioral state and independently of its actual spatial context. The nature of hippocampal network activity during such out-context retrieval has not been described so far, though. Theoretically, context-independent spatial memory retrieval suggests a shift from the hippocampal spatial representations coding the actual- to the remembered context. In this study we show in rats that CA3 neuronal population can switch spontaneously across representations and transiently activate another stored familiar spatial pattern without a direct external sensory cuing. This phenomenon qualitatively differs from the well described sharp wave-related pattern reactivations during immobility. Here it occurred under theta oscillatory state during an active exploration and reflected the preceding experience of sudden environment change. The respective out-context coding spikes appeared later in the theta cycle than the in-context ones. Finally, the experience induced as well an emergence of population vectors with a co-expression of both codes segregated into different phases of the theta cycle.

## Introduction

A prerequisite of episodic memory recall is the ability to reactivate previously encoded neural activity patterns independently of the current spatial context. It is self evident that in humans it can occur across different awake behavioral states such as immobility or active locomotion. Episodic memory in humans as well as spatial memory and allocentric navigation across mammalian species critically require functional hippocampus ^1,2^. Regardless of whether or not we consider the laboratory animals capable of remembering a full set of human episodic memory dimensions – semantic, spatial, temporal and emotional – understanding of how hippocampus encodes and how it recalls the spatial information provides an important knowledge closely related to the mechanisms of episodic memory in general.

Hippocampal principal cell activity is strongly modulated by subject’s position in space, forming context-specific neural representations, that likely provide a physiological substrate of spatial memory^3^. Memory recall in general is a very volatile process between the cue appearance and the behavioral response. When thinking about an episodic memory recall, one would assume that its spatial component requires a reactivation of the respective spatial representation in the hippocampal networks, shifting thus from the code for subject’s actual location to the stored spatial code of the episode that might have occurred in entirely different context. Analysis of spiking activity from populations of individual place cells enables to identify the evolution of spatial information code in a high temporal resolution and to capture eventual transient shifts between in-context and out-context related hippocampal representations.

In the literature on rats and mice, there is a considerable amount of studies on reactivation of hippocampal spatial representations that were encoded in a different context than the one the animal occupies at the time. However, were shown during sleep^4–6^ or during animal’s awake immobility^7^, in both cases when the networks expressed an ‘off-line mode’ characterized by large irregular activity (LIA) with occasional sharp wave/ripple complexes (SWR). These instances are rather considered as part of consolidation process, than the element of memory recollection^8–10^. While the animal performs locomotion, theta oscillation (6-12 Hz) dominates in the hippocampal local field potential and the place cell population activity mainly codes for its actual position. There is a growing amount of knowledge about the ‘non-local’ patterns generated during ongoing theta. They refer to distant positions, but within the same environment context, rather reflecting planned or considered actions to be performed within the imminent future ^11–14^. A spontaneous out-context spatial pattern recall that would emerge during the awake theta oscillatory state was not described so far.

Jezek et al. (2011) showed the hippocampal CA3 network can transiently switch from a representation reflecting the current environment to a map of an independent context and eventually flicker back and forth for couple of seconds, while being paced by the local theta oscillation. The emergence of this phenomenon immediately followed the ‘teleportation procedure’ in which the rat experienced a sudden change between the visual sensory inputs identifying the two respective familiar environments. The transient retrieval of out-context pattern could be as short as a single theta cycle and its scattered emergence lasted only for couple of seconds after the sensory switch^13,15^. Given it’s short life it is likely it originated from the conflict between the visual inputs that responded to the changed cue conditions in the apparatus, and the self-motion input. After the path integrator reset, the network flickering stabilized into the state that corresponded to the actual environment.

In this work we searched for a retrieval of out-context place cell patterns in hippocampal CA3 network that would 1) emerge during ongoing theta activity, and 2) to occur spontaneously without being immediately preceded by any kind of external cuing. Rather, we exposed the animals to the experience of contextual cues switch that, as shown previously, causes the shift in the respective hippocampal representations, and we searched for a spontaneous occurrence of the map shifts after a longer delay. We found spontaneous recalls of the out-context spatial CA3 patterns corresponding to the alternative contexts in sessions as delayed by as 20-60 minutes after the context-switching experience.

## Materials and Methods

### Subjects

14 male Long-Evans rats ranging from 6 to 9 months of age and weighting between 400 – 500 g were housed individually in Plexiglass cages with transparent cover with stable 12 h/12 h light/dark cycle. Experiments were performed during the light phase. In days of training, the animals had limited access to food and were kept around 90% of their free-feeding body weight, while water was accessible ad libitum.

All protocols followed in this study were approved by the Ethical Committee of the Ministry of Education, Youth and Sports of the Czech Republic (approval no. MSMT-10669/2016) according to the Guide for the Care and Use of Laboratory Animals (Protection of Animals from Cruelty Law Act No. 246/92, Czech Republic).

### Electrode preparation and surgery

Rats were implanted with a ‚hyperdrive’ containing circular bundle of 16 independently movable tetrodes. Tetrodes were twisted from four 17 μm polyimide-coated platinum-iridium wires (90% and 10%, respectively; California Fine Wire Company). Their tips were plated with platinum to reduce their impedances to 120–200 kΩ at 1 kHz.

Before surgery, the animals were food deprivated for 12 h. Anesthesia was induced by placing the animal in an enclosed box filled with isoflurane vapor and then the intraperitoneal injection of ketamine (90 mg/kg) and xylazine (10 mg/kg) was delivered. The animal was fixed into a stereotactic apparatus with continuous influx of air (1500 ml/min) containing 1.5-3% isoflurane. The Isoflurane flow was regulated depending on breathing and reflex patterns. The tetrodes were placed above CA1 of the right hippocampus at coordinates of 3.8 mm posterior and 3.00 mm right to bregma. The hyperdrive was then anchored to the skull with jeweler screws and embedded into dental acrylic. Two screws served as electric ground. After the surgery, rats were allowed to recover for one week before the training started.

### Tetrode positioning and recording procedure

Over the course of two weeks following the implantation, tetrodes were slowly lowered towards their intended location in CA3 of hippocampus. Movements proceeded in steps of 50 μm or less and were halted when a large-amplitude theta-modulated complex-spike activity appeared. One tetrode was left in corpus callosum to serve as a reference. Several hours before the experiment started, the signal was fine-tuned. Tetrode’s final positions were checked histologically after the experiment had finished.

The hyperdrive was connected to a multichannel, impedance matching, unity gain headstage. The output of the headstage was conducted via a lightweight multiwire tether cable and 82-channel slip-ring commutator to a data acquisition system containing 64 digitally programmable amplifiers (Neuralynx, Bozeman, MT, USA). Unit activity was amplified by a factor of 3000-5000 and bandpass filtered from 600 to 6000 Hz. Spike waveforms crossing an individually set threshold (30-80 μV) were time-stamped and digitized at 32 kHz. EEG signals, 1 per tetrode, were amplified by a factor of 1000 and recorded continuously (bandpassed between 0.5 and 475 Hz) at sampling rate of 2 kHz. Light emitting diodes (LEDs) on the headstage were used to track the animal’s position every 40 ms.

### Behavioral training procedures and experimental setup

Both experimental and control groups contained 7 subjects, respectively. The procedure used was based on the protocol developed by Jezek et al. ^13^. The training phase was designed to establish orthogonal spatial representations of two environments (identical 60×60 cm square arenas enclosed by 40 cm high black walls on white matte translucent plexiglass floor). Both contexts differed by unique configuration of controllable LED lights on the walls and beneath the floor. The recording area was surrounded by black curtains to eliminate any external visual cues. The arena LEDs were the only available light sources.

To strengthen discrimination between both contexts, subjects received environment-specific food incentives. Before every trial, the arena was thoroughly cleaned with diluted detergent.

The training consisted of four stages (Fig. 1A). In the first stage (double box), arenas were located next to each other and connected by a 20 cm wide and 20 cm long passage allowing the animal to travel freely between the enclosures in three 20 minutes sessions.

**Figure 1.**
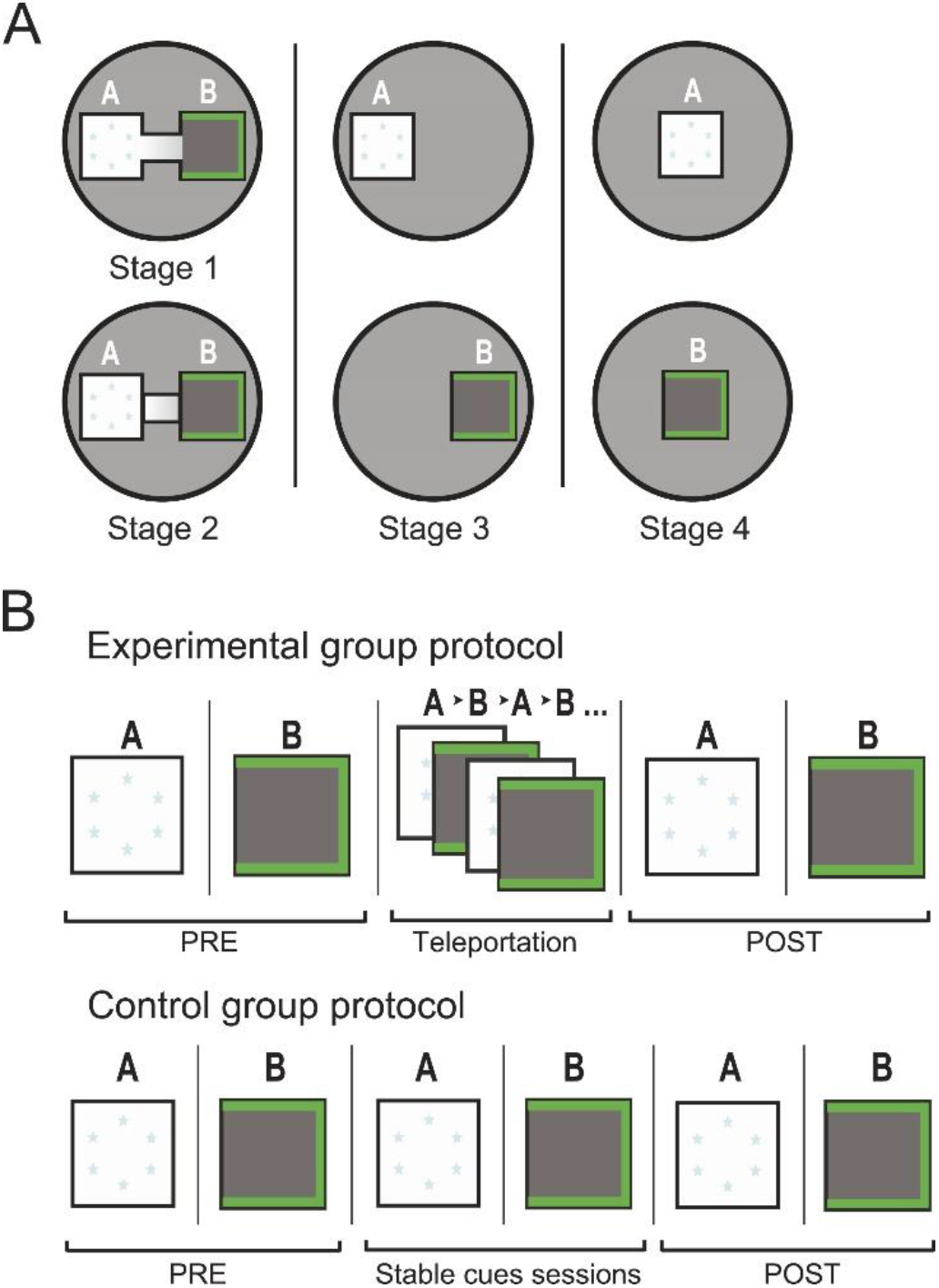
Scheme of the training and experimental paradigm. (A) All procedures were performed in an enclosure surrounded by black curtains. The training scheme was designed so, the animal forms two distinct spatial representations for the respective environments. (B) Scheme of the experimental day procedure. Initial baseline recordings (PRE) were followed with the teleportation session in experimental group or with two sessions with steady cues in controls. After 20 and 50 minutes, respectively, two test sessions (POST) were recorded.

In the second stage (double box disconnected), the connecting alley between arenas was closed and the animals received three 10 min sessions in each of the environment in a semi-random order. In the third phase (single box/two locations) we replaced the whole apparatus with one single box that was equipped with both sets of lights and presented at the original locations. The animals received again three 10 min sessions in each version of the environment. The final forth stage (single box/one location) was based on presenting the single box from the stage 3 in the common location in the center of the recording chamber independently of the cues lit. The rats experienced three sessions in each of the two environment identities. The order of environment presentation across phases 2 to 4 was semi-random. Across all phases the animals rested for 20 minutes between the sessions in a cushioned pot on a pedestal located outside of the curtains.

On the testing day (Fig. 1B), first, both groups were exposed to each environment for 10 min (PRE sessions). Then, the experimental group received one teleportation session. Each rat was introduced into the arena with one set of lights on. After 60-90 s of free exploration the light cues were switched to the other set, instantaneously changing the visual discriminants of the environment. The light switching (teleportation trial) repeated every 60-90 s for 10 min. The control group animals received two 5 min sessions, one per each environment identity spaced with 5 min break, instead of the teleportation session. Afterwards, both groups received again one 10 minutes session in each environment in the stable cue conditions (POST sessions).

### Spike sorting and cell classification

Offline spike sorting was performed manually in three-dimensional feature space depicting projections of waveform amplitudes and energies (SpikeSort 3D 2.5.1.0, Neuralynx). Only well-separated clusters were accepted into the analysis. Putative pyramidal cells were identified by their spike width, average firing rate and by distribution of their interspike intervals (occurrence of complex spike bursts). Putative interneurons were not included into the analysis. Only data from periods with movement velocity larger than 5 cm/s were included in the analysis.

### Template rate maps construction

To define the expected spatial distribution of cell firing, the template population vectors were constructed based on the activity from PRE-sessions 1 and 2. We first split the PRE-sessions in halves and used the first part of each session to establish the activity templates, and the second half for the baseline measures. In both halves the rats sufficiently covered the arena surface.

First, the registered track coordinates were transformed into 20 × 20 position bins (pixels) of size 3 × 3 cm. Then the spatial ratemaps were built by dividing number of spikes emerged in each bin by the total time spent in the given bin. The resulting ratemaps were smoothed using Gaussian boxcar average over the surrounding bins.

### Momentary population vectors

Theta waves were identified from local EEG using a Hamming window and a band-pass filter between 5/6 Hz and 11/12 Hz. The cell activity-based population vectors (PV) were constructed by assigning the individual cell activity into the temporal bins defined by periods of the successive theta cycles. The borders between the bins were set to theta phase with the lowest mean firing rate across the collected cell population ^13^. Each population vector was linked with a corresponding position of the rat derived from the tracking file.

### Population vector categorization

The contextual specificity of place cell activity for each position bin was assessed using ‘Position/Environment Specificity Index’ (PESI), that defined the difference between the cell’s expected activity across the environments for the given position bin. Then, for the position bin x, PESI_x_=(f_A,x_ – f_B,x_) / (f_A,x_ + f_B,x_), where f_A,x_ and f_B,x_ are the reference mean firing rates for the given bin in the respective environments. The place cell activity was considered specific in position x if the corresponding PESI_x_ had value of -1 or 1. The analysis was restricted only to those position bins that showed activity from at least 2 cells with PESI = 1 and another 2 cells with PESI = -1. This ensured that we were capable of registering both context-specific patterns (A and B, respectively) for the given bin whenever they should occur. We considered only spikes from those cells that showed context-specific activity in the given position bin. (Fig. 2A).

**Figure 2.**
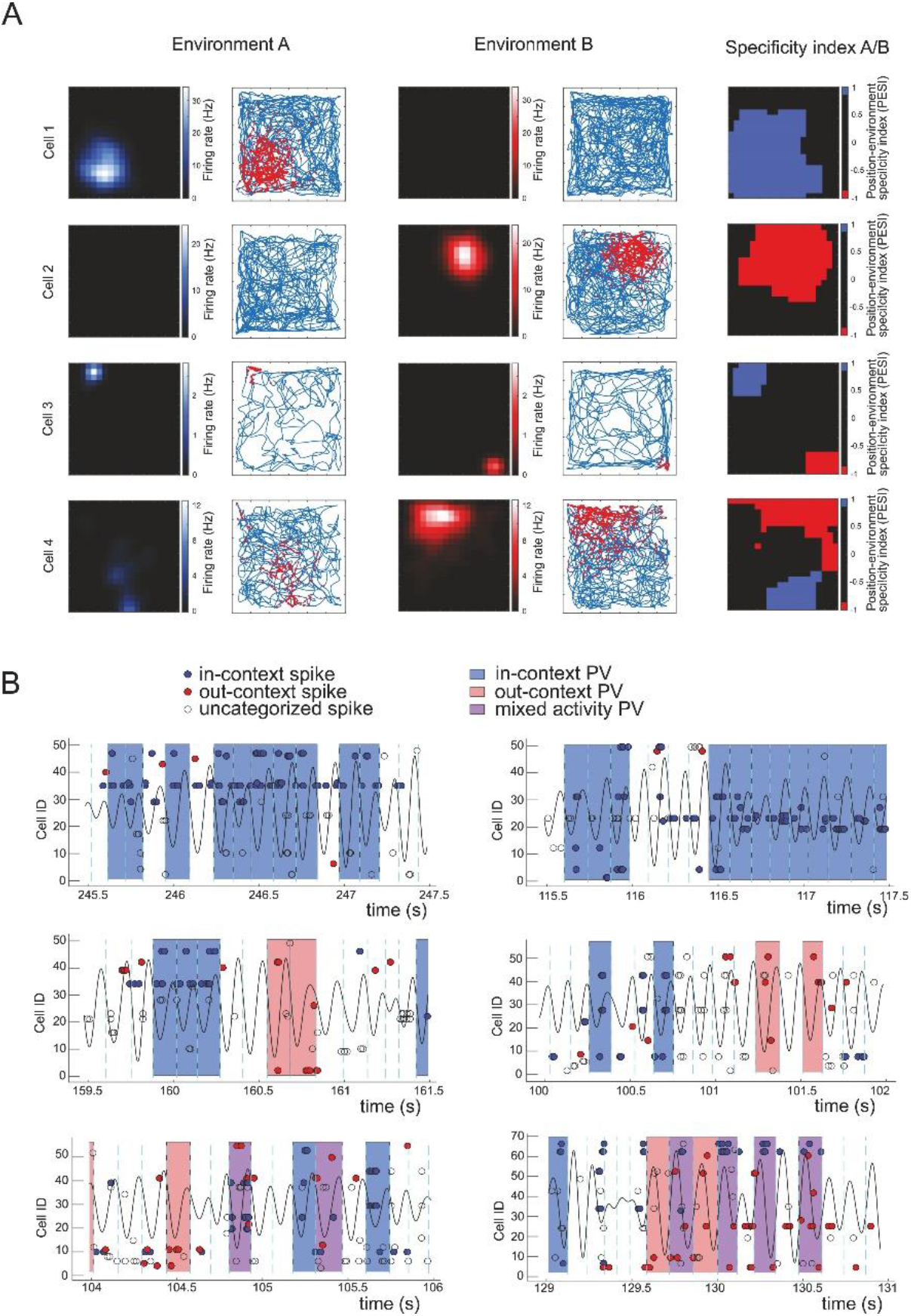
Examples of cell and population activity. (A) Examples of activity from four individual CA3 cells across environments A and B. First four columns represent ratemaps and track (blue line) with superposed spikes (red dots) from template sessions in environments A and B, respectively. The column on the right displays position bins where the cell demonstrated the activity specific to one of the environments, either A (blue) or B (red). Some cells (# 3 and 4) were active in both environments. Only the cell’s activity in bins with the absolute specificity index was considered in later analysis. (B) Examples of population activity evolution and categorization. Population activity patterns categorized as in-context (ICPV) are undercolored with blue, the out-context (OCPV) with red, and the mixed patterns (MPV) with purple. The criteria are described in Methods section. Black bold curve represents filtered LFP (8-12 Hz) and the dashed cyan lines show identified borders between the population vector bins. Spikes (circles) from individual cells (in lines) are marked with red or blue in case they fired at the position with an absolute specificity coding, measured by the cell’s environment specificity index. Spikes that were expected to fire at according the pre-defined template in the environment present at the time, are highlighted in blue. Spikes that were expected to fire at the given coordinate only in the alternative environment are highlighted with red. Spikes categorized as nonspecific (empty circles) appeared in spatial bins without the absolute specificity for any of the environments and were disregarded from PV analysis.

Depending on the cells active in the momentary population vector, were categorized the data into ‘in-context’, out-context’ and ‘mixed’ population vectors. The first pattern, the ‘in-context population vector’ (ICPV), was identified when the given PV contained activity from two or more cells specific to the context present at the moment, and no spike from a cell specific to the alternative environment. The second pattern, the ‘out-context population vector’(OCPV), was scored when the PV contained spikes from 2 and more cells specific to the alternative context at the given position bin, while no spike from the cell specific to the present context. Identification of ‚mixed’ activity pattern (mixed population vector, MPV) required simultaneous activity of at least two cells otherwise specific for each environment in given spatial pixel. As specified above, the only pixels considered were those encoded by sufficient number of specific place cells allowing to detect all activity types.

To control for differences in cumulative times spent across various coordinates, we normalized the activity pattern counts. The amounts of each pattern scored in the given position bin was normalized by the total amount of temporal bins spent there. All corresponding scores were then averaged across the whole session. They are presented in the format of the average number of activity pattern instances per 1000 TC. To control for possible differences in results based on unequal cell-sample sizes across the groups, we made 10 randomly selected subsamplings of place cells in the control group to equal the experimental group sample size. Then we reanalyzed the incidences of various activity patterns and averaged results of all iterations.

### Theta phase of activity patterns

All presented phase values are relative to theta cycle borders determined individually for each recording (0 degrees at the theta cycle border). The respective relative theta phase value was detected for each considered spike in the analyzed PV. In mixed cycles, we treated separately spikes from the place cells specific for the current and alternative environment, respectively. To perform the phase statistics, we used MATLAB Circular Statistics Toolbox (Version 1.21.0.0, Berens) using Watson-Williams Test for equality of means with Bonferroni correction for multiple comparisons.

### Spatial and temporal context of activity patterns

The precision of position coding (position coding error) of the individual theta cycle events was assessed using a correlation decoder, where the momentary population vectors consisting of spike counts produced by recorded place cell population during respective theta cycle were correlated with the template population vectors across all the position bins of the arena.

The decoded position corresponded to the position bin with the highest correlation and the coding error was defined as the distance between decoded and the real position of the animal. The coding error was computed for the in-context and out-context theta cycles, respectively, and averaged across the session.

We evaluated temporal relationships between instances of spatial map instability. We measured average temporal distance between subsequent OCPVs and compared them to average temporal separations measured in 5000 samples containing equivalent number of simulated, randomly distributed OCPV instances. Then we focused on OCPV clustering on shorter timescale. We restricted the analysis to sessions containing at least 12 OCPVs. We created binary arrays, corresponding to theta binning of data. The n-th position within an array yielded 1 if n-th theta bin expressed an OCPVand 0 otherwise. The dot product auto-correlations of the arrays were then computed. The auto-correlation values were then normalized by computing a z-score: z = (C-mean(|Cshuffle|))/std(|Cshuffle|), where C was the observed cross-correlation value and |Cshuffle| was the set of crosscorrelation values in shuffled data. The randomization was performed by randomly shuffling position of the OCPVs within theta bin array. The eligible bins for OCPV replacement had to contain activity of at least two cells and occur at a rat position sufficiently covered by place fields to detect expression of the respective spatial maps. The shuffling was repeated 1000 times for each session.

The z-scored values were used to compute the average auto-correlogram across the sessions. The significance of OCPV cumulation at subsequent theta bins was evaluated for each analyzed session by comparing the auto-correlation value for one theta bin lag with the shuffled data. The one-sided theoretical p-value corresponds to p = (n+1)/1001, where n is number of cases in shuffled data with value equal to or larger than in the real data.

### Statistics

Pre-processed electrophysiological and behavioral data were imported into MATLAB (Version 8.4, MathWorks Inc.) and further processed by custom-written scripts. Statistical analysis was performed using MATLAB, STATISTICA (Version 12, StatSoft Inc.), Excel (Version 14.0.7214.5000, Microsoft Corporation) and R (Version 4.4.1, R Development Core Team). Statistical inference was performed using R statistical software ^16^. Residuals of all models were visually checked for potential bias in model predictions and/or heteroscedasticity ^17^. Data of repeated measurements with approximately homoscedastic and normally-distributed errors (data of speed) were analysed using linear mixed-effects models (LME) with animal identity representing random-effect factor (random intercept), using *nlme* package ^18^. Proportions (relative incidence of theta cycles in given state before multiplying by 1000) were analysed using beta regression with logit link function, and with animal identity representing the random-effect factor, via *mgcv* package ^19^. Comparison of position errors between out-context and in-context states were analysed using generalized mixed-effects models (GLMM) with Gamma-errors distribution and log-link function via *glmmTMB* package ^20^. As the the logit-linked beta regression does not allow occurrence of zeros, and zero-inflated models are unsuitable in our case, data were bounded to ¾ of the minimal non-zero value of given vector ^21^. To robustly evaluate the accuracy of the regression coefficients estimates (ß; in log-odds for beta regression, logarithms for Gamma GLMM and raw values for LME), non-parametric bias-corrected and accelerated (BCa) bootstrap (5000 resamplings ^22^) was used to compute 95% confidence intervals. In the case of mixed states incidence, we used percentile bootstrap due to polymodal distribution of simulated estimates and thus possibly inaccurate bias correction. Whole clusters of correlated data were resampled to preserve within-subject dependency ^23^. In analyses of within-session trends, time bin was treated as numerical variable with presumed linear effect since its categorization increased *Bayesian information criterion* ^24^.

### Histology and electrode positions

Rats were terminated using overdose of pentobarbital and were intracardially perfused with ringer solution followed by 4% formaldehyde. Brains were extracted and stored in formaldehyde to be subsequently frozen and cut into 30 μm coronal sections and stained in cresyl violet. Tetrode tips were located by comparing tissue damage in adjacent sections.

## Results

### Identified place cells

Across all subjects we identified 585 place cells (mean 29.25 ± 3.25 cells per rat). For each animal, we found on average of 253.00 ± 19.39 out of 400 position bins that met our specificity coding criteria. In short, they had to be covered by firing fields of at least 2 place cells that would fire in the given pixel exclusively in one out of the two environments.

### Memory trace dynamics

As previously shown a sudden switch between the cues identifying two distinct familiar environments leads to a shift in the respective neural representations of space in hippocampal CA3 and CA1, followed with a transitory network state instability ^13,15,25^. The spike data were binned into population vectors reflecting the individual theta cycles from parallel-recorded LFP, as the theta oscillation has been shown to pace the hippocampal expression of the spatial code ^13–15,26^. The resulting TC-based population vectors from the sessions preceding (PRE) and following (POST) the teleportation session were then categorized as expressing in-context (ICPV), out-context (OCPV) or mixed (MPV) spatial codes (see Methods; Fig. 2B).

We found the experience of repetitive teleportations led to a significant increase of the incidence of out-context population vectors from PRE to POST stable environment sessions approximately 2.5 times, that is from 1.489 ± 0.587 OCPV/1000TC (PRE) to 3.789 ± 1.301 OCPV/1000TC (POST) (Fig. 3A; ß = 0.901 [95% CI: 0.716, 1.468], p < .0001, beta regression). This contrasted with data from control group exposed to stable environment sessions instead of teleportation that did not show any significant change in OCPV (PRE = 2.009 ± 0.429 OCPV/1000TC; POST: 2.345 ± 0.399 OCPV/1000TC, ß = 0.158 [-0.288, 0.514], p = 0.345, beta regression, after PV size normalization (see the methods); PRE: 3.689 ± 0.968 OCPV/1000TC, POST: 4.193 ± 1.255 OCPV/1000TC; ß = 0.128 [-0.304, 0.333], p = .311, beta regression, without normalization). When comparing to dynamics in control group after equalization of place cell sample (as average number of place cells recorded was higher in control group; control mean n = 33.1, experimental mean n = 26.9), the 2.5 fold increase observed in the experimental was significantly higher than the change by factor of approximately 1.2 observed in the control group (ß = -0.665 [-1.217, - 0.245], p = .025, beta regression). The change in instability following intervention did not prove to be significant in the control group. The size of population vectors were equalized due to variation in place cell sample size across groups. This led to difference in baseline incidence of OCPV during PRE sessions between groups in the original data with complete place cell samples (ß = -1.259 [-2.486, - 0.211], p = .029, beta regression). After we equalized the size of population vectors across the groups, the PRE OCPV incidence became comparable (ß = -0.931 [-1.922, 0.069], p = 0.073, beta regression). The results were confirmed by comparing the relative difference in OCPV incidences in PRE and POST sessions. The OCPV incidence increase by factor of 2.5 observed in the experimental group was significantly higher than the relative increase by factor 1.1 in the control group with full PV size (ß = - 0.757 [-1.330, -0.326], p = .004, beta regression).

**Figure 3.**
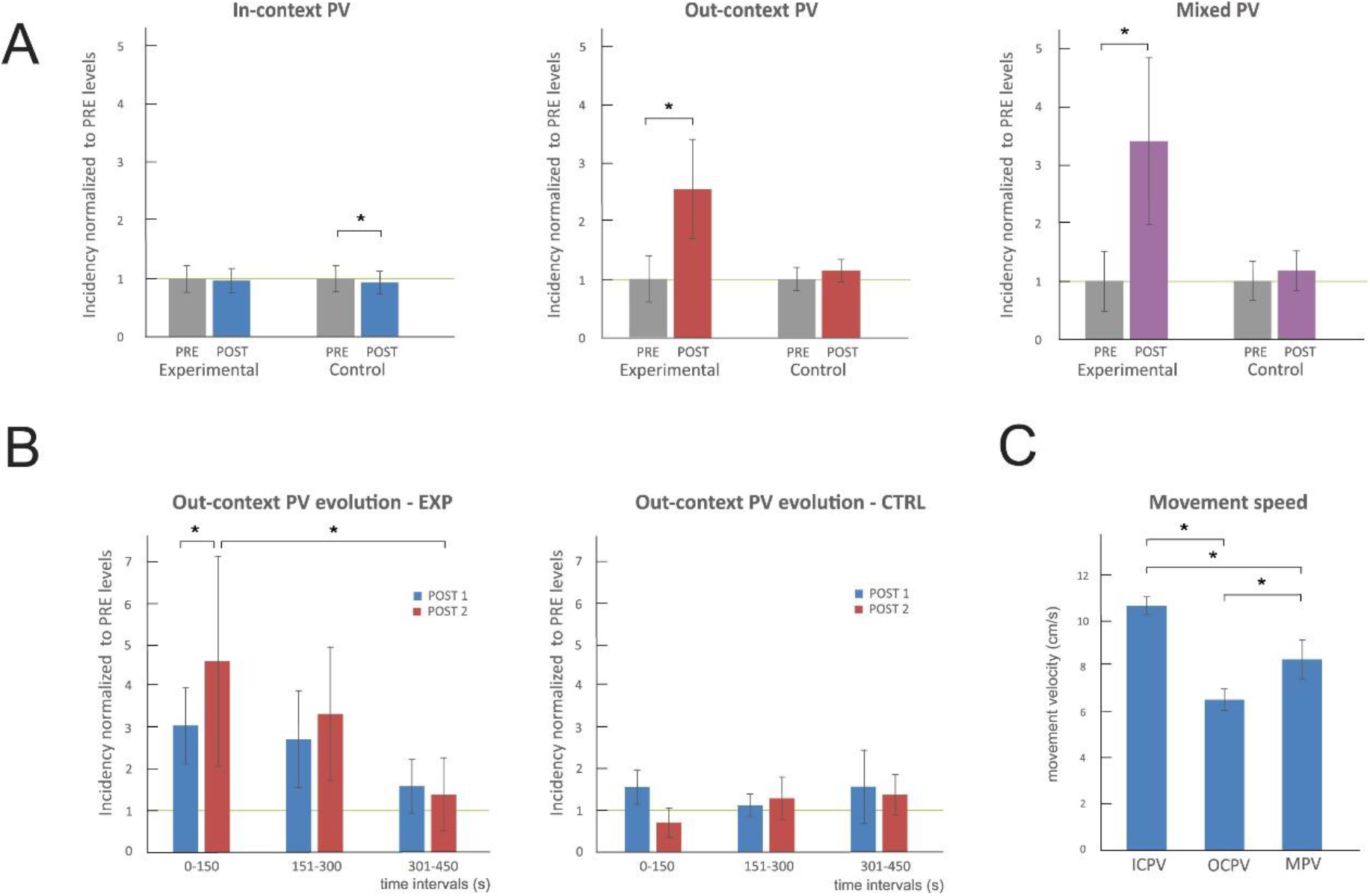
Incidence of in-context, out-context and mixed activity patterns. (A) Relative changes in incidence of all categorized activity patterns across experimental and control group. All values are normalized to the respective PRE-teleportation sessions baseline (green line). (B) Evolution of out-context activity patterns incidence within both POST sessions, divided into three 150 seconds intervals. The emergence of OCPV was high at the beginning of the session and decreased gradually in case of experimental group with the teleportation session experience. (C) Average subject’s movement velocity during each identified activity pattern. * indicates p<0.05.

Similar effect was observed in the mixed population vector analysis (MPV). In the experimental group, the teleportation experience led to an increase in emergence of MPV from 0.639 ± 0.330 MPV/1000TC in PRE sessions to 2.172 ± 1.000 MPV/1000TC in POST sessions (Fig. 3A; ß = 1.099 [0.109, 1.376], p < .001, beta regression). The control group without the teleportation experience did not show such robust destabilization effect and the resulting values were rather comparable (PRE: 0.499 ± 0.163 MPV/1000TC; POST: 0.653 ± 0.186 MPV/1000TC; ß = 0.209 [-0.141, 0.907], p = .408, beta regression, after PV size normalization; PRE: 3.815 ± 1.751 MPV/1000TC, POST: 4.489 ± 1.942 MPV/1000TC, ß = 0.159 [0.109, 1.376], p = 0.055, beta regression, without normalization). The increase in MPV emergence by factor of 3.4 in experimental group was significantly higher than the 1.3 fold increase observed in the normalized control group (ß = -0.990 [-1.279, -0.036], p = .008, beta regression).

Neither the experimental nor the control group demonstrated a change of movement velocity across the sessions. We found the velocity during the correct PV was higher than in the ICPV and MPV, respectively (ICPV:10.72 ± 0.40 cm/s; ICPV 6.59 ± 0.48 cm/s; MPV 8.27 ± 0.88 cm/s). The movement speeds differed significantly from each other (Fig. 3C; ICPV x OCPV: ß = -3.918 [-4.561, -3.098], p < .001; ICPV x MPV: ß = -1.778 [-2.809, -0.930], p < .001; OCPV x MPV: ß = 2.140 [0.999, 3.103], p < .001, all were LME, corrected for multiple testing), but the changes in OCPV and MPV incidence can not be explained by changes in subjects’ activity levels.

The presented destabilization effect was confirmed with alternative criteria requiring only one highly specific cell to fire in the given population vector to be categorized as ICPV, OCPV or MPV, respectively. As above, the incidence of OCPV in the experimental group gained from 21.058 ± 3.293 OCPV/1000TC during PRE to 35.646 ± 6.204 OCPV/1000TC in POST (ß = 0.545 [0.376, 0.691]; p < .001, beta regression). In control group, this effect was not observed (PRE: 29.828 ± 6.559 OCPV/1000TC, POST: 35.192 ± 6.828 OCPV/1000TC, ß = 0.136 [-0.029, 0.301], p = .106, beta regression, after PV size normalization). The increase in OCPV scores was significantly stronger in the experimental group than in controls (ß = 0.428 [0.195, 0.662], p < .001, beta regression). In case of mixed activity, both experimental and control groups displayed a significant increment in MPV. The experimental group showed an increase by factor 1.9 (PRE: 19.445 ± 5.020 MPV/1000TC, POST: 36.315 ± 9.289 MPV/1000TC, ß = 0.659 [0.481, 0.787], p < .0001, beta regression), while by factor of 1.2 in the control group (PRE: 17.207 ± 1.118 MPV/1000TC, POST: 21.267 ± 2.750 MPV/1000TC, ß = 0.208 [0.066, 0.351], p = .004, beta regression, after PV size normalization), respectively. The MPV increment measured in the experimental group was significantly higher than in the control group (ß = 0.453 [0.265, 0.641], p < .0001, beta regression).

In the next step we analyzed the evolution of network instability across the whole sessions. We studied whether network state instability emerged at a constant rate across the set of stable sessions or displayed a mechanistic decrease. In Jezek et al. ^13^, the emergence of OCPV immediately after the teleportation event decreased quickly with time as the network state stabilized. Because in the constant environments sessions the amounts of OCPV and MPV events were much smaller, we divided all PV recorded in the stable sessions (PRE and POST) into three within-session intervals and analyzed the evolution of the respective OCPV emergence. We found a stabilization effect within the POST sessions in the experimental group data. The average incidence of OCPV decreased between the first and the last session’s third from 4.567 ± 1.641 OCPV/1000TC to 1.775 ± 0.706 OCPV/1000TC when both POST sessions were pooled (Fig. 3B; ß (1 time bin) = -0.496 [-0.849, -0.301], p < .0001, beta regression). When analyzing the data from POST1 and POST2 sessions individually, we found a similar trend, however, reaching significance only in the POST2 (POST1: from 3.637 ± 1.137 OCPV/1000TC to 1.894 ± 0.952 OCPV/1000TC, ß = -0.204 [-0.484, 0.012], p = .234, beta regression; POST2: from 5.496 ± 3.143 OCPV/1000TC to 1.657 ± 1.095 OCPV/1000TC, ß = -0.621 [-0.817, -0.435], p < .0001, beta regression). There was significantly higher initial instability in POST2 session than in preceding POST1 session (ß= 0.895 [0.367, 1.241], p < .001, beta regression). The fact the effect was stronger in the second POST session was rather surprising and suggests the network stabilization after a conflicting sensory experience has a long-term non-linear pattern. No such effects of changing instability were observed in the control group (POST from 2.298 ± 0.540 OCPV/1000TC to 2.944 ± 1.058 OCPV/1000TC, ß = -0.008 [-0.270, 0.347], p = .933, beta regression, normalized PV size; from 3.820 ± 1.226 OCPV/1000TC to 4.691 ± 1.888 OCPV/1000TC, ß = 0.067 [-0.209, 0.294], p = .523, beta regression, not normalized PV). None of the PRE sessions neither in the experimental, nor in the control group demonstrated such evolution in within-session OCPV presence (EXP: from 1.112 ± 0.556 OCPV/1000TC to 1.178 ± 0.438 OCPV/1000TC, ß = 0.038 [-0.236, 0.312], p = 0.785, beta regression; CTRL: from 1.761 ± 0.362 OCPV/1000TC to 1.719 ± 0.640 OCPV/1000TC, ß = - 0.008 [-0.195, 0.179], p = .933, beta regression, normalized PV size). The average MPV scores in the experimental group changed from from 2.515 ± 1.251 MPV/1000TC to 1.280 ± 0.623 MPV/1000TC during POST sessions, but their low individual counts were statistically non-evaluable.

### Temporal pattern of OCPV/MPV emergence

We evaluated whether instances of instability of network activity in the form of OCPV expression followed any temporal pattern. The mean interval between the two subsequent OCPV events was significantly shorter than their random distribution across the whole session (data: 46.511 ± 4.923 s; random: 71.018 ± 7.020 s; U(n_1_=n_2_=63) = 3487, z = - 2.503, p = .012). Because this difference might not only point to the within-session restabilization effect shown above, we focused closely on the short inter-event intervals. The analysis was restricted to a subset of sessions (n=21) showing at least 12 OCPV occurrences. We constructed binary PV arrays mapping the emergence of OCPV across the given sessions, respectively (Fig. 4A), and calculated their autocorrelation. We found a considerable cumulation of short lag values peaking at 1 theta bin lag (Fig. 4B; one-sided p-values: p < 0.05 in 18/26 sessions). This suggests that while majority of detected OCPV occurred in temporal isolation (Suppl. Fig. 3), there was relatively frequent emergence of out-context patterns within subsequent theta cycles.

**Figure 4.**
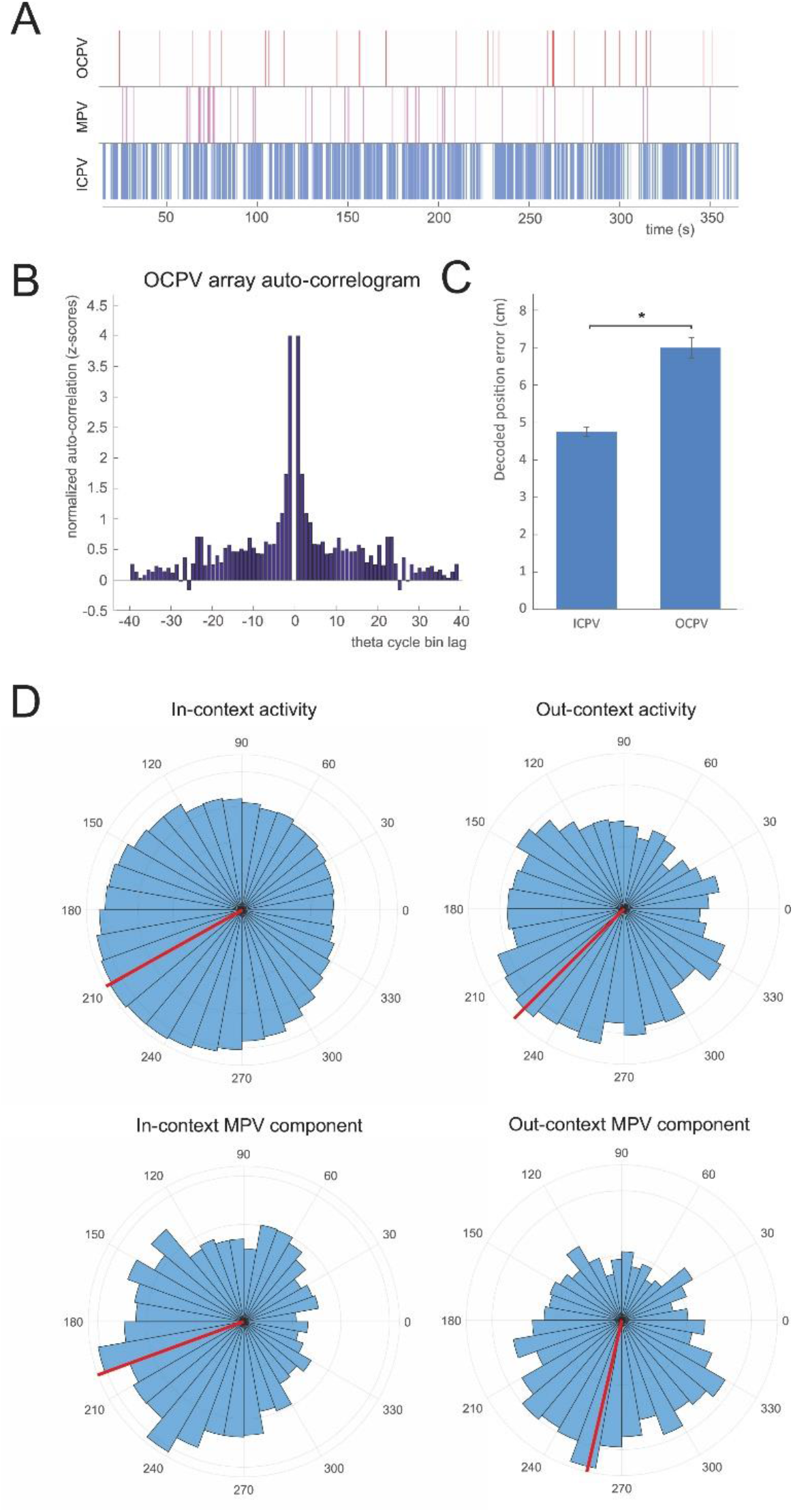
Properties of the identified activity patterns. (A) Example of temporal distribution of classified activity patterns during one of the POST sessions from an experimental group animal. Top: out-context PV, middle: mixed PV, bottom: in-context PV. le MPV and red OCPV; vertical separation of activity types serves for better visual differentiation of activity instances. (B) Out-context PV incidence auto-correlogram. Values were z-scored using random distribution and then averaged across sessions. (C) Deviation between the decoded and actual animal’s position across OCPV and ICPV. Out-context PV events demonstrated higher position error than ICPV ones. (D) Distribution (blue) and average (red) theta phase of spikes from identified in-context, out-context, and mixed population vectors. Phase values are normalized to the individual identified phase border between the theta bins (see the Methods section). Spikes from OCPV appeared in significantly later phases of the theta cycle than ICPV. The in-context and out-context activity components of the mixed PVs appeared in significantly different portions of the theta cycle. * indicates p<0.05.

To evaluate quality of spatial code during reactivation of the alternative representation, we applied spatial position decoder on activity vectors corresponding to OCPVs. The positional error was higher when decoding from activity during OCPV than during control in-context states (ICPV: 4.765 ± 0.122 cm, OCPV: 7.021 ± 0.274 cm, ß = 0.366 [0.283, 0.449], p < .001, Gamma GLMM).

### Theta phase shift

In the next step we checked the relation of identified population vector patterns to local theta oscillation. All the analyzed PVs were associated with animal locomotion and majority of them occurred under strong theta oscillation. Momentary power of theta oscillations relative to the rest of the oscillatory spectrum did not differ between PV patterns (ICPV 13.204 ± 0.443, OCPV 12.279 ± 0.462, MPV 12.032 ± 0.725; F(2, 186) = 1.49, p = 0.228).

Then we analyzed the theta phase locking for spikes occurring in the correctly, incorrectly and mixed coding population vectors. To deal with the unequal number of categorized theta cycles in different animals, we first compared the respective average phases obtained across the individual subjects. We found the OCPV spikes occurred on average 27.14° later than those from ICPV cycles (ICPV: 205.93 ± 4.28°, OCPV: 233.07 ± 6.62°; F(1,146) = 9.358, p = .026). In case of mixed population vectors, we split their content into in-context and out-context activity (in relation to the actually present environment) components based on the same criteria used for the PV assignment. On average, the in-context spikes preceded the out-context ones by phase shift of 86.94° within the given theta wave (in-context: 210.22 ± 8.85°, out-context: 297.16 ± 9.81°, F(1, 80) = 24.558, p < .001).

This result was confirmed by comparison of all averaged phases taken from the individual population vectors of the same type pooled together irrespecitvely of the subject. The respective values for ICPV (209.28 ± 0.15°; Fig. 4D) were significantly different from spike phases belonging to OCPV (225.00 ± 0.97°; F(1, 255123) = 86.92, p < .001; Fig. 4D). Large phase shift was confirmed between in-context (200.14 ± .24°; Fig. 4D) and out-context (257.05 ± 1.47°; Fig. 4D) components of the mixed PVs (F(1, 6007) = 318.17, p < .001).

## Discussion

In this paper, we described occurrence of spontaneous transient recall of hippocampal CA3 place cell patterns referring to a context that was different than the one the subject was currently actively exploring. We showed that the emergence of out-context patterns increased after the experience of repetitive sudden contextual changes delivered by the teleportation procedure^13^ and it tended to stabilize across the post-teleportation stable-cues sessions. We found that spikes of the two pattern classes (the in-context and the out-context), respectively, showed a different theta phase modulation. The cell activity coding the regular in-context pattern appeared earlier within the theta cycle than those from the pattern referring to the alternative environment context.

In recent years and decades, various forms of non-local coding have been described under different conditions – some refer to more or less distant positions within the actually present context^11,12,14^, others to entirely different one. The expression of population code referring to true remote context has been iconically connected to the neural pattern replays during sharp wave/ripple events. This phenomenon is considered to be part of memory consolidation^8–10^. Its occurrence is usually observed in a resting box, where the animal dwells immobile or sleeping right after the experience in the experimental apparatus, or during transient immobility epochs of active exploration^7^. In such situations the hippocampus expresses large irregular activity dominated by slow waves and the circuit is to a large extent disconnected from sensory information flow as if it was in an ‘off-line mode’.

In contrast, during active exploration the hippocampal local field potential is dominated by theta oscillation (8-12 Hz) and the populations of place cells mostly code for animal’s actual position. Besides this, there is growing body of evidence that the considered or planned actions are reflected during the theta state as well, in a form of activating the place cell patterns that correspond to more or less distant locations or position code sweeps withing the present context. Hippocampal place cell sweeps were initially described in the vicarious trial and error paradigm (VTE) ^14^, when the rat was preparing for the left or right turn response at the choice point of the continuous T-maze. While ongoing theta rhythm, subsets of place cells that corresponded to the available routes from the choice point were activated in a sequential manner ‘modelling’ the future responses in a time-compressed manner. More recently, a constantly present theta sequences ^27^ were identified as the animal was either actively exploring the environment or preparing for a behavioral choice. Both in the cases of VTE and various theta sequences observations the animal generated ‘mental sweeps’ into possible locations within the same environment context that might represent a neuronal substrate of planning the immediate future responses.

In contrast, this report shows emergence of patterns related to entirely different context while exploring the environment under the ongoing theta oscillation. The described spatial representation shifts were inserted within periods of the actual position coding, and were rather sparse and short, ranging from a single to several theta cycles. They resemble the theta paced flickering described in ^13,15^ in response to sudden contextual change induced by the teleportation protocol. However, theta paced flickering phenomenon followed immediately the substantial sensory change in form of contextual shift and lasted only for several seconds before the network stabilized in the activity state corresponding to the newly present context. This nature suggests the flickering ^13^ might be rather a result of competition between the idiothetic and allothetic information inputs (path integrator vs. visual cues) resulting in rapid shifts between the respective attractor states^28^ or a short-term plasticity^29^. The instances of activating the out-context pattern here, however, occur under substantially different conditions. They come during the sessions with steady context present, dozens of minutes after the last teleportation event. Such conditions turn the possibility that the out-context pattern expression rises from the conflict between the hippocampal inputs as rather unlikely.

From point of attractor network theories the hippocampal representations of space behave as continuous attractor states of neural activity, tolerating subtle deviations of their inputs from the stored activity patterns^30–33^. Whereas in the case of sweeps and theta sequences the non-local population activity travels within the same continuous attractor state, the phenomenon presented here showed a transient network activity shifts into a different manifold in absence of any related sensory stimulus. This suggests that their emergence was driven intrinsically. While their amount was minimal in the first sessions, the teleportation experience led to their significant increase during the sessions that followed. However, the experimental evidence that would point to a mechanism leading to a sensory-independent shift between the representations is missing. A possible explanation would be that the preceding sequence of teleportation events with the respective representation shifts brought both representations into a temporal proximity that might reinforce an associative binding between the two. Such a link then might have increased the probability of a spontaneous retrieval of the alternative context pattern during the sessions that followed.

However, this explanation somewhat contrasts with another interesting observation, which is the progressive reduction of the out-context pattern events rates during both POST stable-context sessions. This state-change dynamics was indeed induced by the prior teleportation procedure, as we did not see any such development in the control group. Owing to the seen progression, the out-context pattern recall was rather temporary and tended to settle down as the session was going on. It is worth noting that this stabilization effect did reset after the first post-teleportation session because it replicated itself in the following one with comparable magnitude. This suggests, that a factor of expectancy of another teleportation trial might contribute to the out-context patters activation at the beginning of each post-teleportation session.

We noted as well spontaneous appearances of mixed patterns between the representations for both contexts within the individual theta cycle temporal bins. Again, this observation had its analogy to what has been described immediately after the teleportation trials during the supposed conflict between the path integration and visual information inputs^15^. Here, however, the mixed patterns emerged during stable context cue sessions taking place long time after the teleportation session, and in a limited amount even before it. Moreover, we observed a phase locking structure between the spikes specific to the present and alternative context within the mixed PV bins, respectively. The cells firing specifically in the currently present environment tended to appear earlier in the theta cycle compared to cells that were active exclusively in the alternative context. This within-theta organization resembles recently reported constant theta based referencing to possible futures in hippocampus^12^ where the neural activity alternates between the code for present location and the possible future path, respectively. That phenomenon shows a similar phase locking properties with the mixed activity introduced in this report, as in Kay et al. (2020) the information about the present location also preceded the estimated future within the given theta cycle. From this point of view, the ‘mixed patterns’ might reflect referencing to the alternative environment context rather than the non-organized mixture of the two context representations, because the out-context part of the code tended to follow after the in-context portion of the mixed pattern. The phenomenon is experience driven as the instances of the out-context retrieval were more frequent after the repetitive teleportation protocol that suddenly changed the context identity and induced the respective representation shift. However, such a rate increase was rather temporary, and the network tended to stabilize within the behavioral sessions. In this respect the out-context pattern retrieval shares the key features with the concept of episodic memory retrieval, namely in the context-independency and by its spontaneous and transient occurrence. The existence of episodic memory in laboratory animals is a matter of long-going debate stimulated by numerous experiments in rodents or other species^34–36^. We speculate that the out-context pattern emergence might offer a valuable model for network mechanisms of episodic memory recall. The transitory and spontaneous nature of out-context and mixed PV emergence as well points towards the instability of the network activity states that represent a broadly discussed substrate of some psychiatric disorders, namely the manic disorder and schizophrenia. In this relation the reactivation of context-unrelated network activity patterns might account for the emergence of positive symptoms such as delusions or hallucinations^37,38^.

