## Supplementary figures for "Context-independent expression of spatial code in hippocampus"

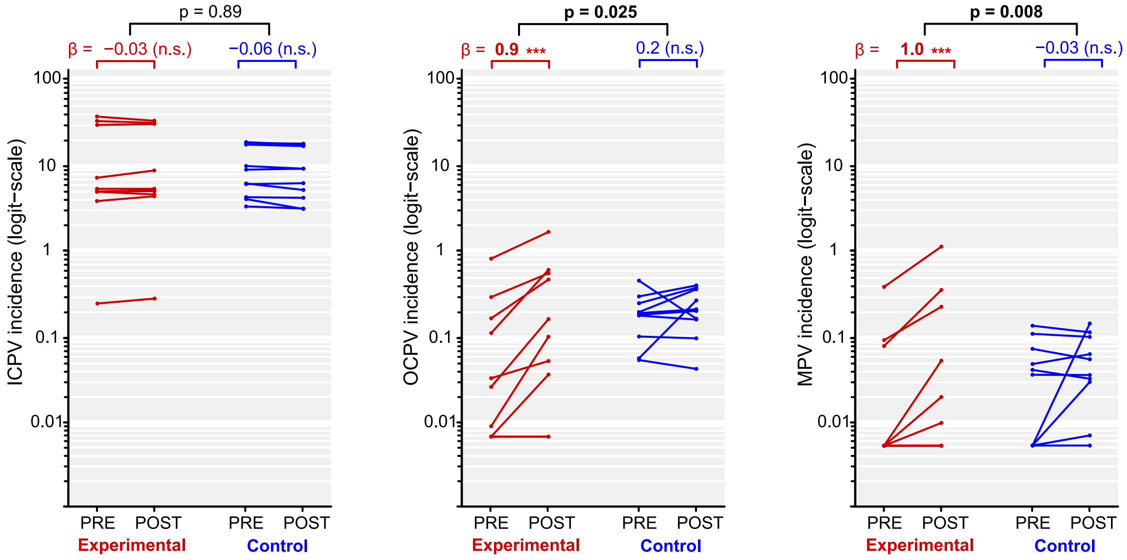


**Supplementary figure 1. Changes of incidence in the identified network activity patterns across individual experimental days.**

Left panel: In-context PV; midle panel: Out-context PV; right panel: mixed PVs

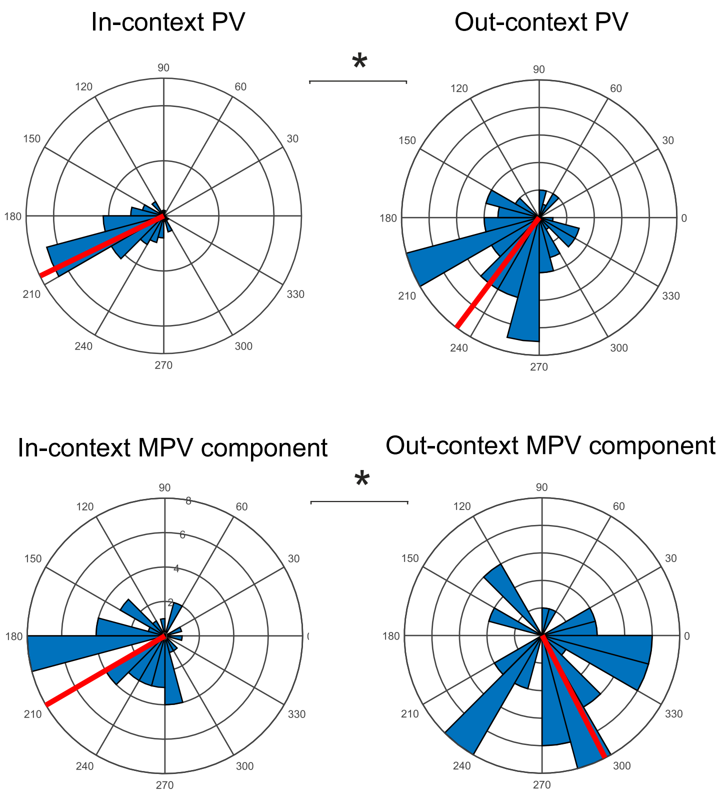


**Supplementary figure 2. Distribution of average spike theta phase locking across individual experimental days.**

Top left: In-context PV; top right: Out-context PV; bottom left: in-context spikes from mixed PVs; bottom right: out-context spikes form MPVs. Phase values are normalized to the individual identified phase border between the theta bins (see the Methods section). * indicates $p<0.05$.

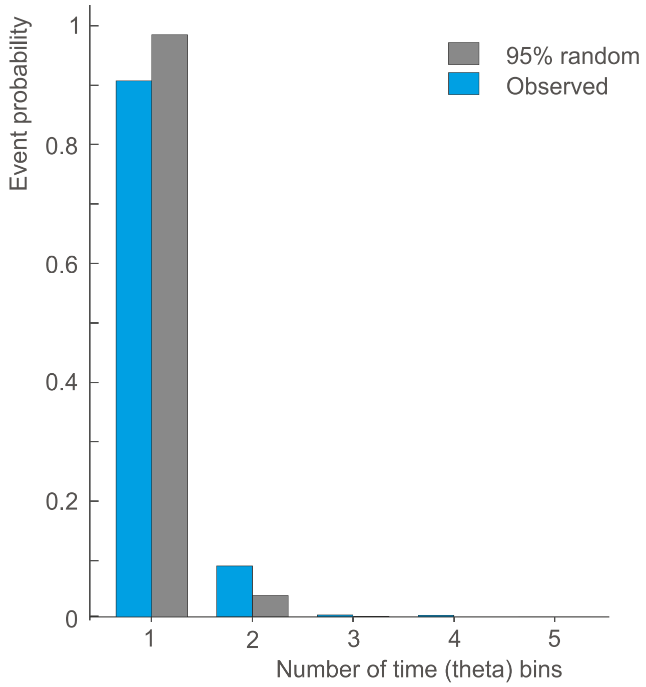


**Supplementary figure 3. Length of the recalled out-context population activity patterns.**

Blue bars indicate the counts of all identified out-context events according their length measured by amount of consecutive time bins. The grey bars show the 95th percentile of shuffled distributions, generated by 1000 times randomly choosing equal number of ICPV. The observed amounts of out-context events longer than 2 and more theta cycle bins significantly exceed their expected random occurence.
